# Design of a Modular Inducible Circuit for Pulsatile Population Control of Engineered Bacteria

**DOI:** 10.1101/2025.05.03.652077

**Authors:** Rohita Roy, Michaëlle N. Mayalu

## Abstract

Synthetic biology enables the precise control of bacterial population dynamics through engineered genetic circuits, paving the way for innovative therapeutic and biosensing applications. Here, we present genetic circuit design that leverages AHL (acyl-homoserine lactone)-based quorum sensing and a transcriptional cascade regulated by an external inducer to drive synchronized proliferation and lysis events. In this system, the absence of the inducer arabinose prevents CinR mediated repression, enabling activation of a transcriptional cascade that drives the expression of the lysis protein, fluorescent protein, and a drug resistance gene. The accumulation of AHL acts as a proxy for population density and modulates gene expression through a quorum-sensing mechanism. This inducible circuit provides tunable and pulsatile control over bacterial population dynamics, enabling cycles of population growth and collapse. Such behavior prevents premature population clearance while ensuring functional bacterial densities for extended durations. Our system represents a foundational step toward implementing more complex multi-stable architectures for self-regulation of bacterial population. This dynamic and modular framework offers significant advantages for therapeutic interventions, such as self-regulating drug delivery, inflammation-responsive probiotic therapies, and targeted bacterial therapy.

## Introduction

The advent of synthetic biology has transformed our ability to engineer living systems by designing and implementing synthetic genetic circuits that reprogram cellular behavior with high precision. These circuits, composed of modular biological parts enable cells to process information, make decisions, and execute complex functions in response to internal and external stimuli.^1,2^ Among the many transformative applications of synthetic biology, the precise regulation of microbial population dynamics has emerged as a critical area of innovation. By designing circuits that modulate bacterial growth, survival, and programmed cell death, researchers can exert fine-grained control over microbial communities—an ability that is essential for the development of advanced microbial therapies, diagnostic biosensors, and environmentally responsive biocontrol systems.^3–5^ Population-level control is particularly vital in therapeutic contexts, where engineered microbes are deployed to colonize host tissues, deliver drugs, or modulate the host immune response. Without proper control, such microbes may overgrow, trigger immune clearance, or lose therapeutic efficacy due to stochastic fluctuations in gene expression or evolutionary instability.^6,7^ Genetic circuits that incorporate mechanisms like quorum sensing, inducible lysis, or toxin-antitoxin systems can dynamically regulate population size and function, thereby enhancing safety and effectiveness.^8,9^ In environmental applications, population control circuits are instrumental in bioremediation efforts where bacteria are used to degrade pollutants, as well as in agriculture to prevent the uncontrolled spread of engineered strains.^10^ A major challenge in the development of microbial-based therapies and biosensing platforms lies in achieving a careful balance between sustained bacterial activity and robust biocontainment. Engineered microbes are increasingly deployed in complex and dynamic environments such as the gastrointestinal tract,^6^ solid tumors,^11–13^ where they are expected to detect specific cues, deliver therapeutic payloads.^12^ However, without precise control mechanisms, these microbes may overgrow or persist in unintended locations, leading to potential adverse outcomes. Over proliferation can result in reduced therapeutic efficacy due to immune recognition or resource competition, and may even lead to safety concerns such as sepsis and inflammation.^14^ Conversely, overly stringent biocontainment strategies—such as constitutive kill switches or nutrient auxotrophy—can prematurely eliminate the engineered population before it accomplished its intended function.

This can be particularly problematic in therapeutic applications where continuous or periodic bacterial activity is required over extended durations.^15^ For example, in tumor-targeting applications, bacteria must colonize the tumor microenvironment, reach a therapeutic density, and release cytotoxic agents over time. If containment is activated too early, the bacteria may be cleared before a sufficient therapeutic effect is achieved.^12,16^ Thus, a key requirement for next-generation synthetic microbial systems is the ability to modulate population density dynamically— enabling engineered bacteria to grow, carry out their programmed function, and then self-clear in a temporally regulated manner. This type of control can be achieved through synthetic gene circuits that integrate environmental signals, quorum sensing, inducible promoters, and feedback loops to coordinate population-wide responses.^17–21^ Oscillatory population dynamics, in particular, offer a compelling framework by allowing cycles of expansion and synchronized lysis, thereby sustaining functional populations while minimizing long-term persistence or overgrowth.^9^ Achieving this level of spatiotemporal precision will be critical for the safe and effective translation of synthetic microbes into clinical applications. Traditional approaches to bacterial population control in synthetic biology have often relied on constitutive expression of toxic genes or single-feedback loop mechanisms to limit bacterial growth.^6,12^ While these systems can effectively restrict population size or enforce cell death under defined conditions, they are typically rigid in their response profile exhibiting binary behavior—either fully active or completely repressed—with minimal sensitivity to dynamic environmental cues. As a result, such static designs often fail to sustain viable bacterial populations for extended periods, limiting their applicability in therapeutic or biosensing contexts where persistence, adaptability, and responsiveness are critical.^10^ Furthermore, these simplistic regulatory strategies are ill-suited for complex environments like the gastrointestinal tract, tumor microenvironments, or natural ecosystems, where microbial populations must adapt in real-time to fluctuating chemical, physical, and biological inputs.^4^

To address these limitations, recent advances in synthetic biology have emphasized the development of genetic circuits with dynamic and tunable behaviors. A notable example is the implementation of oscillatory circuits, which enable bacterial populations to alternate between periods of growth and programmed cell death.^12^ These systems often leverage quorum sensing, environmental inputs, and delayed feedback to coordinate community-level dynamics. Quorum sensing, in particular, has emerged as a powerful and evolutionarily conserved mechanism by which bacteria sense and respond to changes in population density through the accumulation of small diffusible molecules such as acyl-homoserine lactones (AHLs)^15,22^. When coupled with synthetic regulatory modules controlling survival or lysis genes, quorum sensing can drive pulsatile dynamics—cycles of expansion followed by synchronized lysis—that maintain functional population levels without uncontrolled overgrowth.^3^ In this study, we construct a dual positive-negative feedback circuit that generates pulsatile bacterial population dynamics by integrating an AHL-mediated quorum sensing population control unit with an arabinose-inducible repressor system (Figure 1). This configuration inhibits population growth in the presence of both arabinose and drugs, and, upon arabinose removal, establishes a steady-state population below the carrying capacity by balancing quorum-induced lysis protein expression with quorum-induced drug resistance gene expression. This design allows user-controlled toggling of population behavior through external addition of input signals, while quorum sensing provides a means of autonomous regulation of cell density. This paradoxical (positive-negative) feedback loop architecture captures key features of dynamic regulation and serves as a foundation for developing autonomous feedback control systems that exhibit multi-stable population dynamics. ^8,19-21^

**Figure 1.**
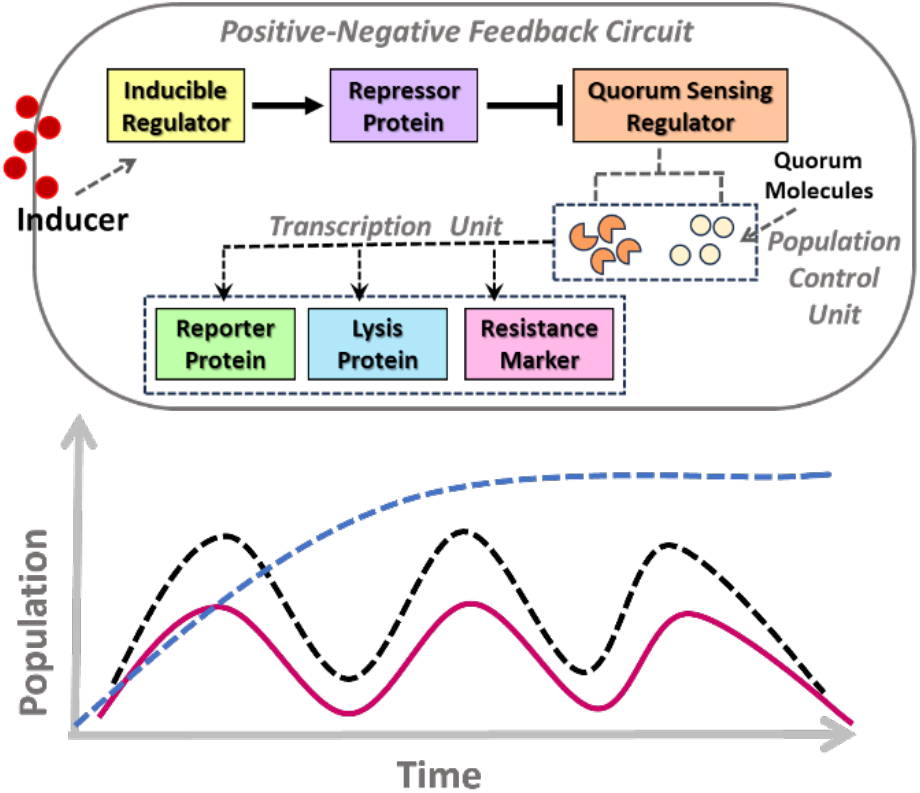
Schematic of a Paradoxical (Positive-Negative) Feedback Circuit for Pulsatile Population Control in Engineered Bacteria. The synthetic circuit integrates an inducible regulator, a repressor protein, and a quorum sensing regulator to modulate gene expression dynamically. The transcription unit encodes a reporter protein, a lysis protein, and an drug resistance marker. External induction triggers the circuit, leading to feedback-controlled population dynamics via a population control unit that senses and responds to cell density through quorum sensing.

The use of the lysis protein offers a potent and well-characterized mechanism for inducing population collapse. Derived from bacteriophage ΦX174, the gene product disrupts bacterial cell membranes, leading to rapid lysis upon induction. When expression is synchronized across the population via quorum sensing, it results in a coordinated die-off of cells (negative feedback), followed by potential regrowth from residual survivors. This rhythmic cycle of growth and lysis mimics naturally occurring pulsatile behaviors and enables sustained population oscillations without long-term persistence.

The inclusion of a drug resistance gene provides an additional, complementary mechanism for regulating population survival. Upon expression, the gene product confers protection against environmental drugs, allowing bacterial cells to survive (positive feedback) under selective pressure. As with the lysis module, synchronization of drug resistance expression through quorum sensing ensures that survival is coordinated across the population. This coupling between drug resistance and cell density enables additional control over steady-state population levels and oscillatory behavior, positioning drug resistance as an active regulator of population dynamics. Importantly, the modular nature of the system ensures its adaptability to different bacterial hosts and quorum sensing systems. The genetic parts used in this system are designed to be standardized, enabling modular recombination or substitution with components from other quorum sensing systems such as LasI/LasR or RhlI/RhlR.^3,23,24^ This modularity also allows for adaptation and optimization in therapeutically relevant bacterial hosts, including *Lactobacillus reuteri, Bacteroides fragilis*, and environmental chassis like *Pseudomonas putida*.^5,25,26^ This versatility enables applications across a wide range of scenarios, including gut-resident probiotic therapies, tumor-colonizing biotherapeutics, and living biosensors designed to detect and respond to environmental cues or disease biomarkers.

One of the most promising applications of this circuit lies in the development of self-regulating drug delivery systems. ^12^ This pulsatile delivery approach offers key advantages: localized drug release, reduced systemic toxicity, and precise control over therapeutic dosing. While prior studies have demonstrated the use of *E. coli* engineered with quorum-sensing lysis circuits for in vivo tumor colonization and pulsatile drug release, these systems often rely on fixed regulatory logic and lack the flexibility to independently tune expression, lysis, and population dynamics.^12,16^ By enabling cyclical control over microbial population growth and clearance, this system lays the groundwork for a new class of intelligent, responsive microbial therapies that can autonomously adapt to changing physiological or environmental conditions.

## Results

### Design and Characterization of the Transcription Unit Synthetic Circuit

Our design offers a uniquely modular and tunable genetic architecture that integrates inducible gene regulation (*via* arabinose and CI), quorum sensing (via CinI/CinR and AHL), and synchronized lysis (via ΦX174) to produce robust, pulsatile bacterial population dynamics. To validate individual components of the paradoxical feedback circuit, we first constructed and tested three strains, each expressing a distinct module under the control of the P_cin_ promoter activated by CinR in response to CinI-AHL. To initiate the design and characterization of our system, we first constructed and tested individual transcriptional units to evaluate the circuit’s performance across a range of CinI-AHL concentrations (10000, 5000, 1000, 100, 10, 1 nM). These simplified modules were designed to independently assess output behavior—including fluorescence intensity and bacterial growth dynamics—in response to CinR–AHL-mediated activation. We quantified gene expression using sfGFP fluorescence (*Strain S1*) **(Figure 2A)** and monitored population growth via optical density measurements at 600 nm (OD_600_). In addition to the reporter module, we also constructed and characterized separate lysis bearing X174 lysis gene protein (*Strain S2*) **(Figure 2B)** and drug resistance modules (*Strain S3*) **(Figure 2C)**, each under the control of the CinR–AHL inducible promoter, to observe their impact on population behavior. These results confirm the successful construction and function of each circuit module, laying the foundation for later integration into the full paradoxical feedback architecture.

**Figure 2.**
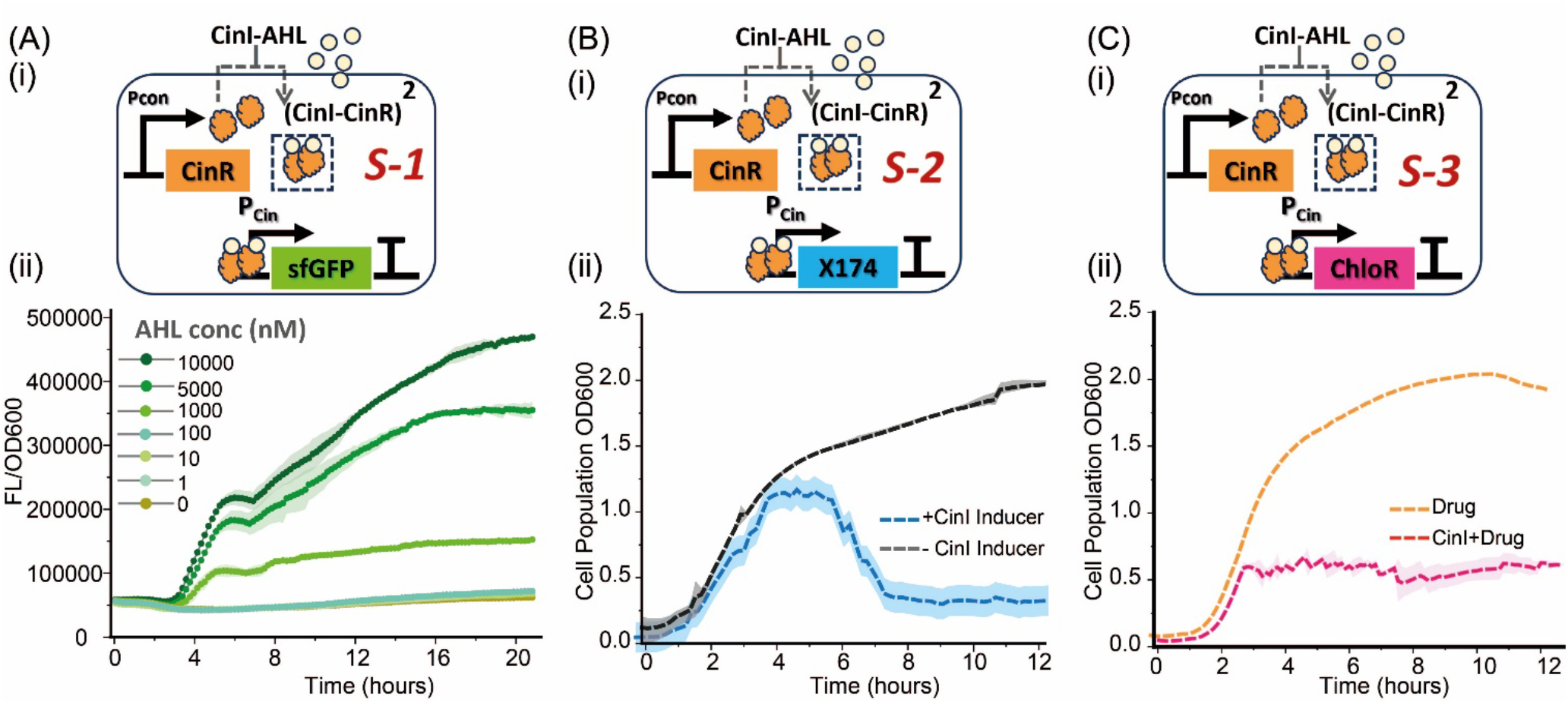
Schematic overview of individual single gene transcriptional circuits. **(A-i)** illustrates construct ***S-1***, in which the expression of sfGFP is driven by the CinR–AHL responsive promoter (P_Cin_) downstream of CinR expression. **(A-ii)** shows the fluorescence response of the circuit across six AHL concentrations (1 nM to 10,000 nM). Fluorescence increases with AHL concentration, demonstrating a tunable and dose-dependent gene expression profile. **(B-i)** shows construct ***S-2***, where the ΦX174 lysis gene is placed under the same P_Cin_ promoter. **(B-ii)** displays cell growth curves with and without CinI induction. Upon induction, lysis is evident as a sharp drop in OD_600_ after ∼4 to 6 hours, confirming effective population collapse triggered by AHL accumulation. **(C-i)** presents construct ***S-3***, incorporating ChloR, a chloramphenicol resistance gene, under same promoter. **(C-ii)** compares cell growth in the presence of chloramphenicol, with and without CinI–AHL induction.

### Steady State Dynamics of Multi-gene Single Input Open Loop Circuit

Following the validation of individual circuit components (*S1–S3*), we next engineered a multi-gene circuit (*S4*) integrating CinI-AHL-inducible expression of sfGFP, ΦX174 lysis protein, and chloramphenicol resistance **(Figure 3A)**. To evaluate the baseline functionality of this composite circuit, we first characterized its steady-state behavior under varying concentrations of CinI-AHL (10000, 5000, and 1000 nM) in both the absence and presence of drug selection (**Figure 3B–E**). At high CinI-AHL concentrations, increased formation of the (CinI–CinR)^2^ transcriptional activator complex led to enhanced activation of the ΦX174 lysis gene, resulting in pronounced cell lysis **(Figure 3B)**.

**Figure 3.**
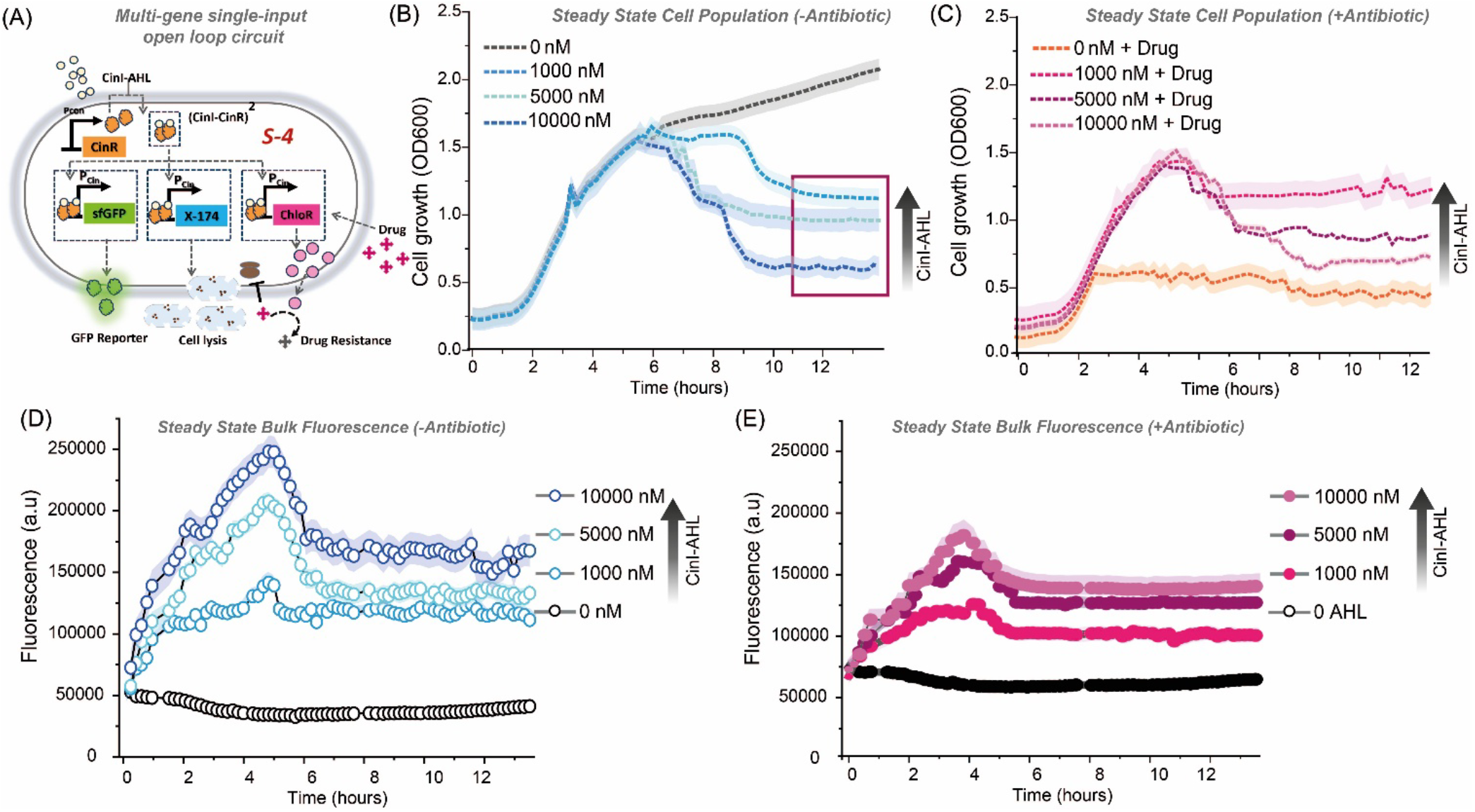
Steady-state characterization of the multi-gene single input open loop circuit (*S4*). **(A)** Schematic of the *S4* genetic circuit architecture. **(B)** Cell growth (OD600) over time for *S4* cultures induced with varying concentrations of CinI-AHL (0, 1,000, 5,000, and 10,000 nM). **(C)** Cell growth (OD600) under the same AHL concentrations in the presence of chloramphenicol (20X concentration drug). Bulk fluorescence output of GFP in the **(D)** absence (open circles) and **(E)** presence (closed circles) of drug across the same AHL concentrations. Fluorescence increases in a dose-dependent manner, indicating successful activation.

In contrast, the addition of chloramphenicol attenuated this reduction in cell density, likely due to its inhibitory effect on ribosomal function,^27^ which may limit translation of the lysis gene and thereby reduce the extent of cell lysis **(Figure 3C)**. Fluorescence output reflects CinI-AHL dose-dependent activation of the genetic circuit and is modulated by drug presence. In the absence of drug (**Figure 3D**), sfGFP expression exhibited a clear, dose-dependent increase in bulk fluorescence following CinI-AHL induction, with the highest signal observed at 10,000 nM AHL. This response confirms robust activation of the P_cin_ promoter and successful signal transmission through the CinI–CinR system. The fluorescence peaked within the first few hours post-induction, followed by a gradual decline likely attributable to lysis-induced population reduction. These results demonstrate that the circuit exhibits stable steady-state responses over the tested conditions.

### Pulsating Dynamics of Single Input Multi-gene Open Loop Circuit

To investigate the dynamic behavior of the full genetic circuit, we monitored both cell population growth and fluorescence output under varying AHL concentrations and in the presence or absence of antibiotic drug selection. **Figure 4** illustrates the emergence of pulsatile dynamics, characterized by cyclic rises and falls in both OD_600_ (cell density) and sfGFP fluorescence over a 24-hour period. To maintain the experimental fidelity after each lysis event we manually removed lysed cells from the culture by centrifugation, followed by multiple washes (wash time points indicated by arrows in Figure 4) with fresh media to eliminate residual lysis proteins released into the supernatant. The surviving cell population was then resuspended in growth fresh medium, enabling continued monitoring of dynamic behavior. This lysis-clearance step was performed twice during the course of the experiment, as detailed in the Methods section. The **Figure 4A-B**, highlight the overall behavior across high AHL concentrations (10,000 nM), revealing a clear pattern of stepwise population collapse and regrowth, probably coinciding with oscillatory gene expression. Drug treatment (20X chloramphenicol) amplifies the effect by imposing an additional selective bottleneck. In the absence of AHL, cells exhibit continuous growth and minimal fluorescence. The moderate AHL condition (5000 nM), the system still exhibits robust pulsatile behavior, although with slightly reduced oscillation amplitude. The fluorescence output continues to follow the same cyclical pattern, supporting the synchronization of gene expression with population dynamics **(Figure 4C-D)**. The lowest AHL condition (1000 nM), results in dampened or irregular oscillations, consistent with a reduced quorum signal input. Here, the presence of the drug slightly lowers the baseline population and fluorescence levels, but the pulsatile pattern remains detectable, albeit weaker. **(Figure 4E-F)**.

**Figure 4.**
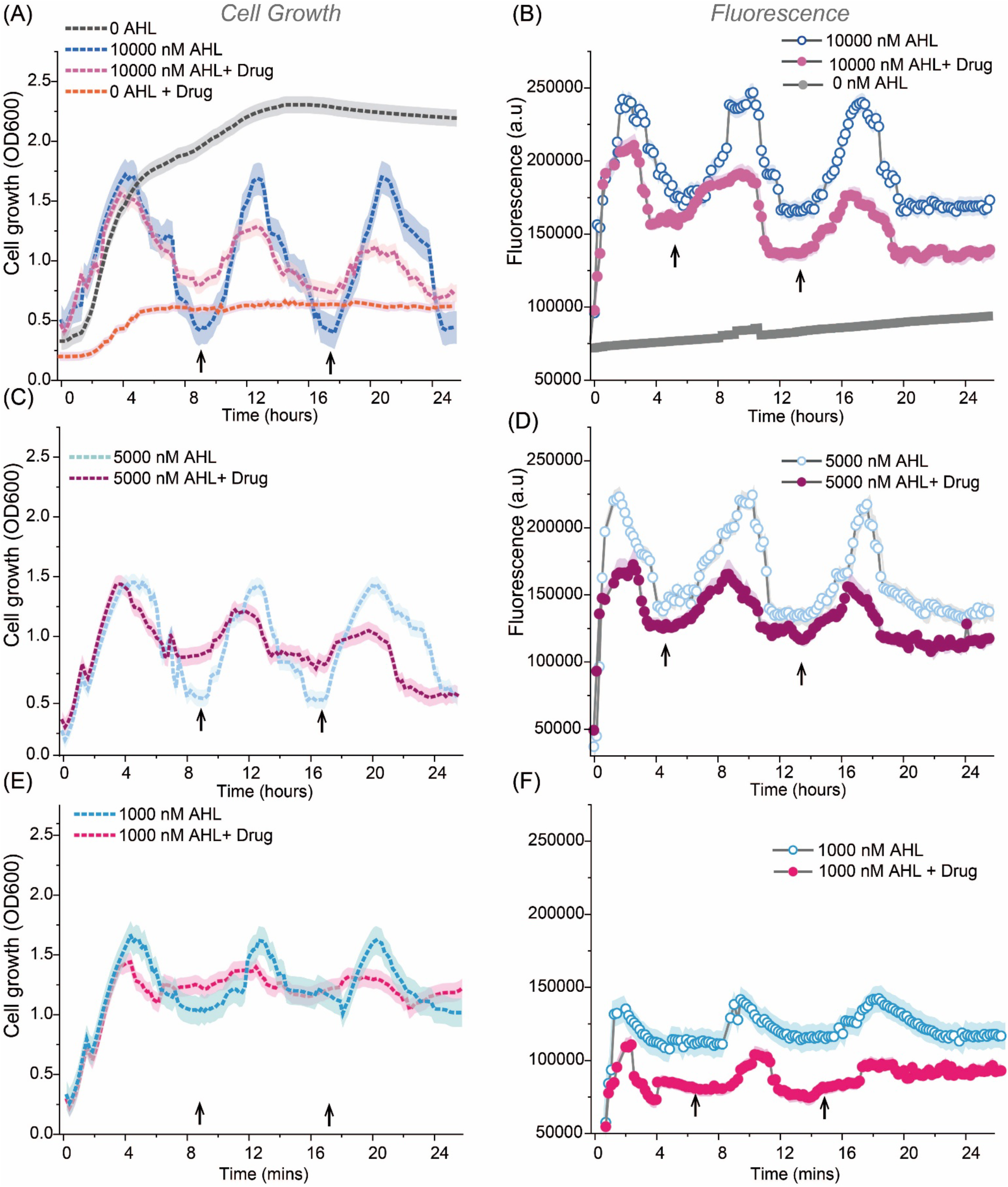
Pulsatile population dynamics of single-input multi-gene loop circuit in presence of drug under varying AHL concentrations. **(A, C, E)** Cell growth measured as OD_600_ over time at high (10000 nM), intermediate (5,000 nM), and low (1,000 nM) AHL concentrations, respectively, with and without antibiotic. Stepwise collapses and regrowth phases indicate rhythmic population behavior at higher AHL levels. **(B, D, F)** Corresponding fluorescence measurements (a.u.) showing oscillatory sfGFP expression that aligns with population growth cycles. Pulsatile dynamics are prominent at high and intermediate AHL concentrations and dampened under low AHL. No oscillations are observed in the absence of AHL or under drug-only conditions. Arrows suggest the time-points of cell washing and introducing fresh media.

### Population Dynamics Regulation in Multi-gene Dual Input Open Loop Circuit

To further realize autonomous, cyclical population control, we constructed a genetic circuit *(S-5)* that integrates environmental sensing (arabinose), and external quorum sensing molecules. The designed system architecture operates under two modes. In the presence of arabinose **(Figure 5A-i)** the AraC transcriptional activator drives expression of the CI repressor, which suppresses CinR and thereby inhibits activation of downstream genes (sfGFP, ΦX174 lysis, and ChloR), keeping the circuit OFF. In the absence of arabinose **(Fig. 5A-ii)**, CI repression is relieved, enabling CinR expression. Upon reaching quorum (*via* CinI–CinR complex formation), the system activates expression of the effector genes, triggering both lysis and drug resistance, and initiating pulsatile dynamics. We tested seven different input conditions **(Figure 5B–C)** combining presence or absence of arabinose, CinI AHL, and 20X chloramphenicol drug, to map steady-state behaviours of the circuit. Under non-inducing conditions in the absence of drugs (states i–iii), cells grew normally with low background fluorescence. When only AHL was present (state iv), and repression by arabinose and drugs was absent, the (CinI-CinR)^2^ complex formed, activating the circuit and inducing strong fluorescence followed by eventual lysis. When drug selection was added without AHL (states v, viii), growth was impaired due to lack of resistance, even in the presence of CinR production. State (vii), characterized by the absence of arabinose and presence of AHL and drugs, exhibited initial growth in the presence of 20X antibiotic, followed by lysis—indicating activation of the transcriptional machinery. Fluorescence studies validated the cell growth dynamics (**Figure 5 D, E**). As shown in **Figure 5F**, the complete induction condition generated robust **pulsatile population dynamics**, with repeated cycles of growth and collapse, confirming that paradoxical feedback enables dynamic homeostasis via integrated control of lysis and survival. In the presence of antibiotic, similar oscillatory behavior was observed but with reduced amplitude, suggesting that the metabolic burden imposed by the drug slows down the pulsation frequency and dampens the overall system dynamics.

**Figure 5.**
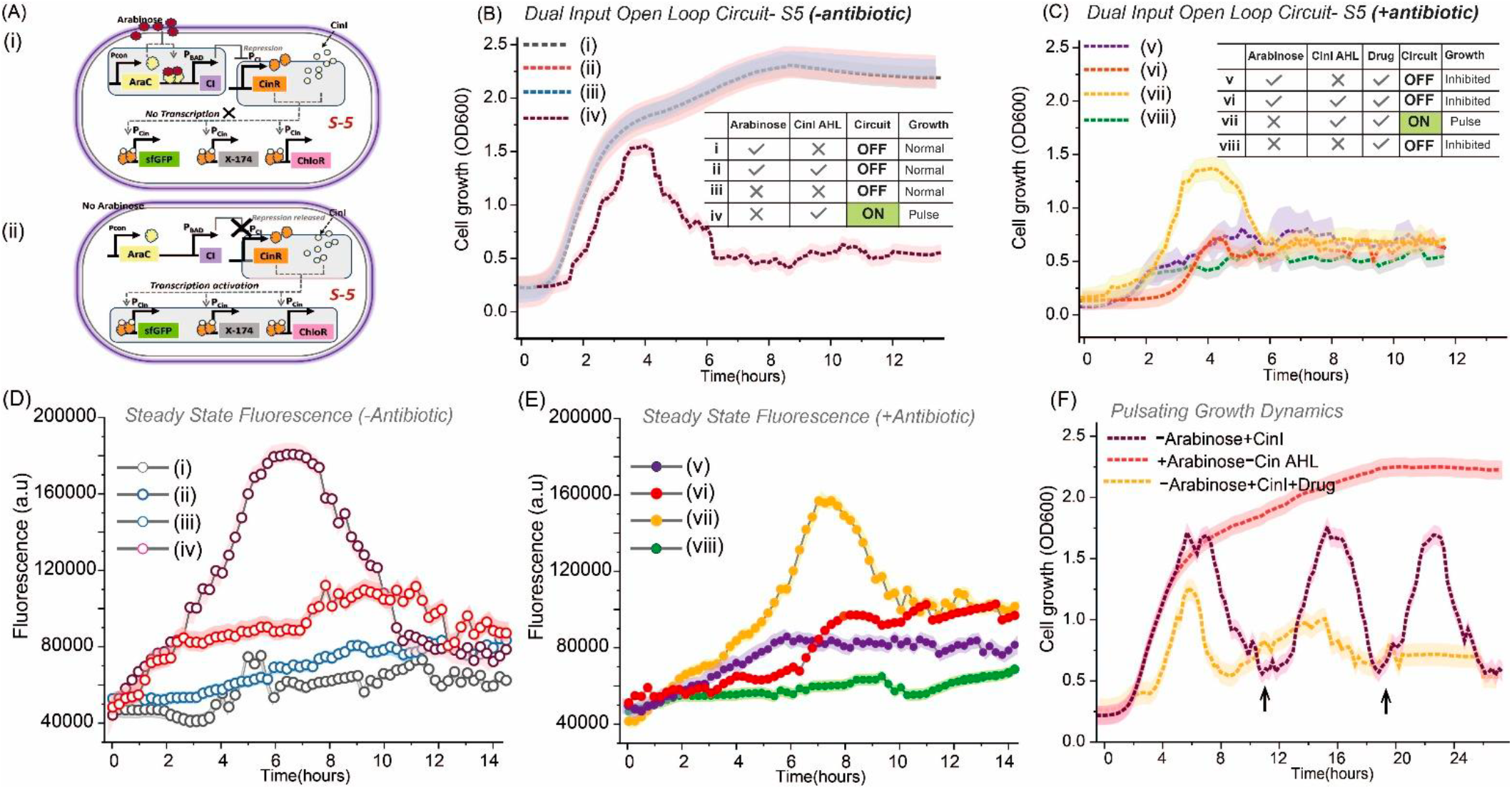
Characterization of the dual input open-loop genetic circuit *(S-5)* **(A)** Schematic of circuit *S-5* under **(i)** repressed (arabinose present) and **(ii)** active (arabinose absent) states. AraC activates CI, which represses CinR and downstream expression from P_Cin_. Removal of arabinose relieves CI-mediated repression, enabling AHL–CinR activation of sfGFP, ΦX174 (lysis), and ChloR (antibiotic resistance gene). **(B-C)** Growth curves under different steady-state input combinations (labeled i–iv) for absence of antibiotic and presence of antibiotic (labelled v-viii). Inset tables summarizes conditions and resulting phenotypes. **(D-E)** Corresponding sfGFP fluorescence outputs for steady-state condition in absence **(D)** and presence of drug **(E)**. Circuit activation **(iv and vii)** yields strong fluorescence. **(F)** Pulsatile population dynamics observed under full circuit activation (+CinI,Arabinose, +CinI and Drug), showing rhythmic cycles of growth and collapse indicating effective paradoxical feedback control. (Arrows suggest the wash time points)

### Population Dynamics of the Closed Loop AHL-feedback circuit driven with Arabinose Regulation

Building upon the *S-5* architecture, we designed circuit *S-6*, which incorporates endogenous AHL synthesis to create a fully autonomous paradoxical feedback loop. While *S-5* required external addition of AHL for circuit activation, *S-6* introduces constitutive expression of CinI (AHL synthase) under the control of a separate constitutive promoter, enabling the bacterial population to self-generate the quorum sensing signal. The results suggest upon removal of arabinose, CI expression is halted, allowing CinR production, followed by circuit activation and activation of the transcriptional cascade **(Figure 6A-B)**. When the circuit was monitored under undisturbed growth conditions for ∼24 hours (**Figure 6B**), it exhibited spontaneous, self-sustained pulsatile dynamics, characterized by periodic cycles of population expansion and collapse. Notably, while the first pulse was prominent, the second pulse displayed reduced amplitude—suggesting partial damping of the oscillatory behavior, potentially due to delayed quorum re-activation or cellular adaptation. A similar pattern was observed under drug selection pressure (chloramphenicol), though the second pulse was markedly attenuated, likely due to the additional metabolic burden imposed by drug resistance expression and drug stress. These findings highlight the circuit’s capacity for autonomous rhythmic behaviour, driven by the integrated control of survival and lysis, and underscore the influence of metabolic load on dynamic output amplitude.

**Figure 6.**
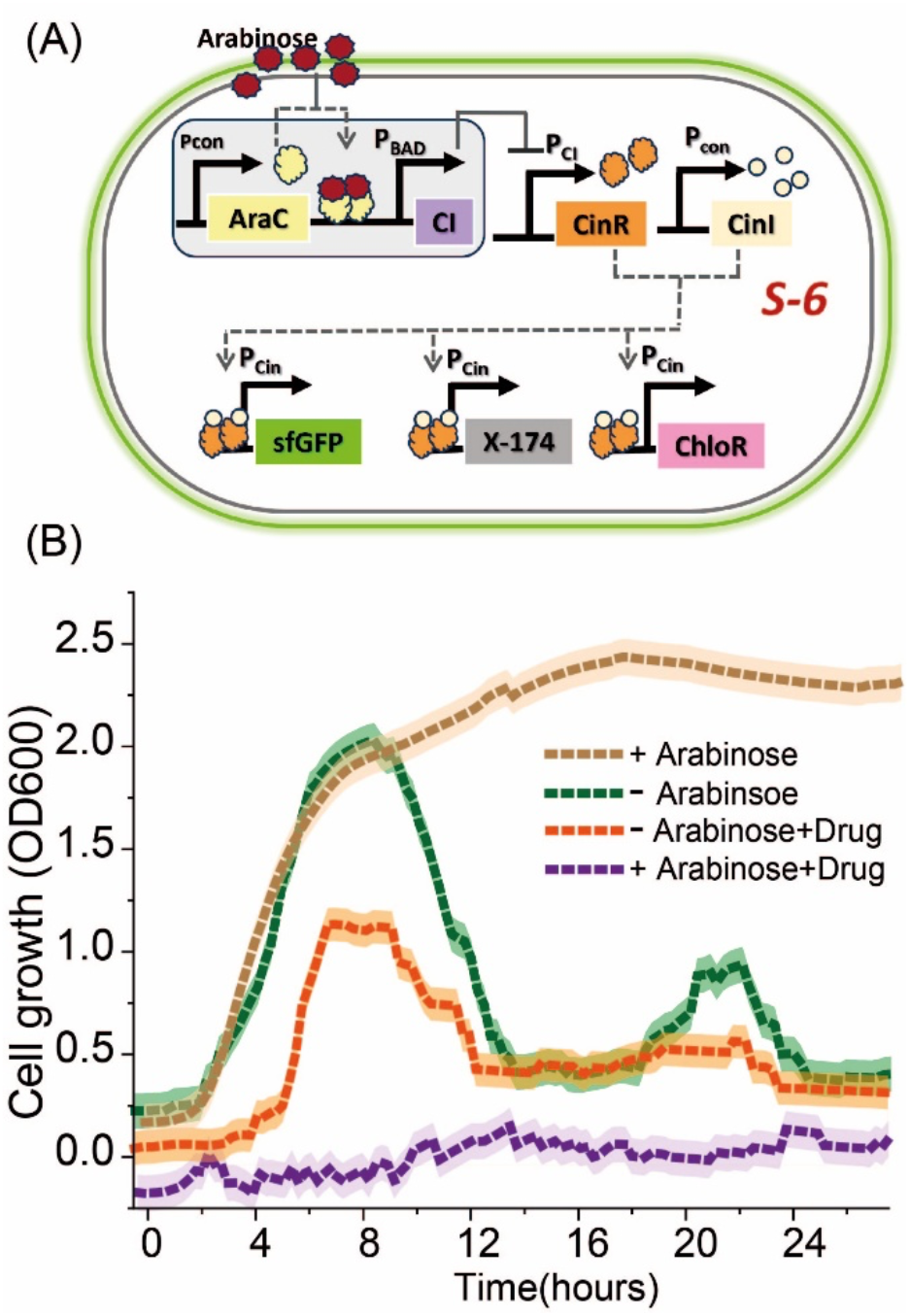
Autonomous paradoxical closed loop feedback circuit (S-6) with endogenous quorum sensing signal production. (A) Circuit schematic showing arabinose-mediated repression and internal AHL synthesis. Absence of arabinose activates the circuit. (B) Cell growth curves showing continuous growth under repressive conditions (+ arabinose), pulsatile dynamics upon de-repression (−arabinose), and sharper, dampened oscillations when chloramphenicol is included (– arabinose + drug), validating the self-regulating, paradoxical behaviour of the closed system.

## DISCUSSION

Together, these results validate a novel genetic circuit that enables inducible, pulsatile population dynamics through the integration of quorum sensing, CinR-mediated population repression, and synchronized lysis. The open-loop design allows users to externally initiate or halt circuit function via arabinose, while AHL modulates internal behavior in a density-dependent manner. Importantly, we demonstrated that the timing, amplitude, and persistence of population pulses can be tuned via AHL concentration, and that drug stress further shapes pulse patterns—either amplifying or attenuating them depending on the quorum threshold. This dynamic flexibility is highly desirable in real-world applications. For example, in microbial therapeutics, pulsatile behavior enables bacteria to accumulate at a site of interest—such as a tumor or inflamed tissue—before initiating drug release via lysis. The self-limiting nature of this approach reduces the risk of immune overactivation or systemic side effects. In biosensing applications, discrete pulse signals provide a high signal-to-noise ratio, facilitating accurate detection of transient environmental cues. The genetic circuit S-6 represents a significant advancement toward fully autonomous, self-regulating microbial systems by incorporating endogenous AHL production into a paradoxical feedback loop. Unlike open-loop designs that rely on external inducers, S-6 enables internal quorum signal synthesis via constitutive CinI expression, closing the regulatory loop and allowing the population to modulate its own behavior in response to environmental inputs and cell density. Critically, this circuit exemplifies the core principles of paradoxical feedback, wherein a single signal—AHL—simultaneously promotes survival (via ChloR expression) and induces cell death (via ΦX174 lysis protein). This antagonistic dual-output control creates the conditions necessary for homeostatic regulation: population growth triggers its own collapse, which then allows for regrowth, establishing a self-sustained oscillatory cycle. While further optimization is needed to stabilize long-term oscillations and reduce amplitude dampening, thus design lays a foundation for building more complex circuit designs.

## CONCLUSION

In this work, we engineered and characterized a modular genetic circuit that enables autonomous, pulsatile control of bacterial population dynamics through a combination of inducible gene regulation, quorum sensing, and paradoxical feedback. Building from an open-loop architecture to a fully closed-loop design, we demonstrated that endogenous AHL production coupled with arabinose-mediated repression can drive synchronized cycles of bacterial growth and lysis. These oscillations arise from the paradoxical role of AHL, which simultaneously promotes survival via drug resistance and induces population collapse via lysis gene expression. Our results highlight the tunability and robustness of this system, showing that oscillation amplitude and timing can be modulated through environmental inputs like arabinose and drug stress. Notably, the fully autonomous behavior of engineered bacteria establishes a foundation for intelligent, self-regulating microbial platforms. This dynamic circuit design offers broad utility across synthetic biology applications, including targeted drug delivery, inflammation-responsive probiotics, and biosensing platforms, where spatiotemporal control and environmental adaptability are critical. By integrating multiple control layers into a single coherent framework, this work advances the development of next-generation synthetic microbial systems capable of safe, programmable, and adaptive behavior in complex setting.

## METHODS

### Genetic Circuit Assembly

All genetic components used in this study were sourced from the Addgene CIDAR MoClo kit, and assembled using a two-step method involving Golden Gate assembly followed by Gibson assembly. Golden Gate cloning was first performed to construct individual transcriptional units (TUs) using Type IIS restriction enzymes and T4 ligase, as previously described.^28^ The transcriptional units (TUs) were constructed using standardized overhangs and assembled following the CIDAR MoClo design principles. The resulting Golden Gate products were amplified using TU-specific adaptor primers. PCR products were purified using the Zymo Gel Extraction Kit. Purified TUs were then assembled into the linearized V35m vector backbone using Gibson Assembly, following the protocol outlined by Gibson *et al*.^29^ Equimolar concentrations of DNA fragments were used for each reaction, and concentrations were quantified using a Thermo Scientific NanoDrop™ spectrophotometer. The Cin-I gene used in construct S6 was amplified using the primer sequences (Fwd 5’-GCA TCG TCT CAT CGG TCT CAT ATG ATG TTC GTT ATC ATT CAG GCA-3’ and Rev 5’-ATG CCG TCT CAG GTC TAG GAT CTG CCA TCT CCA GGA ATT GG-3’,using the standard phusion PCR protocol, followed by PCR product gel extraction.

### Transformation and Validation

Constructs were transformed into either chemically competent *E. coli* DH5α and *E. coli* MG1655 cells and plated on LB-agar containing 50 µg/mL kanamycin. The constructs *S1-S4* were transformed into marionette cell line MG1655 and *S5-S6* were transformed into *E. coli* DH5α. Plates were incubated overnight at 37°C. Individual colonies were cultured, and plasmid DNA was isolated using the Qiagen Miniprep Kit. The integrity and sequence of each construct were validated via sequencing.

### Plate Reader Experiments

Growth and fluorescence measurements were recorded using the BioTek Synergy H1 Plate Reader under the following conditions: Temperature Setpoint: 37°C, Preheat enabled; Absorbance: Endpoint mode, Wavelength: 600 nm; Fluorescence: Endpoint mode, Filter Set: Excitation 479 nm, Emission 520 nm, Light Source: Xenon Flash, Lamp Energy: High, Dynamic Range: Extended. For both absorbance and fluorescence modes, the read speed is normal and delay was 100 ms

### Cell Culture and Experimental Setup

Validated plasmids were transformed into experimental E. coli strains and grown in Terrific Broth with 50 µg/mL kanamycin at 37°C and 220 rpm overnight. The next day, overnight cultures were diluted into fresh media to a starting OD_600_ of 0.2. A volume of 190 µL of this diluted culture was loaded into a 96-well flat-bottom microplate (Corning) for kinetic measurements. Cells were allowed to grow until they reached mid-log phase (OD_600_ ≈ 0.4), at which point CinI-AHL 3-hydroxytetradecanoyl-homoserine lactone (OHC14) ≥96% purity from Millipore Sigma (51481) was added at the desired final concentrations, by diluting in 100 % DMSO. Additional inducers or antibiotics e.g., arabinose, l-arabinose (Ara) ≥99% purity from Millipore Sigma (A3256), chloramphenicol (≥98% (HPLC), Sigma-Aldrich, C0378) were introduced depending on the specific experimental condition.

### Pulsating Experiment

For strains *S1–S5*, long-term pulsatile dynamics were evaluated over a 24-hour period. Following the onset of lysis, cultures were manually removed from the 96-well plate, washed twice with 1X phosphate-buffered saline (PBS) to remove residual toxins and lysed debris, and subsequently resuspended in fresh Terrific Broth (TB). The resuspended cultures were then returned to identical plate reader conditions for continued monitoring. In contrast, for strain *S6*, experiments were conducted under continuous, undisturbed conditions without removal or washing steps, allowing autonomous circuit-driven oscillations to be observed in real time over the entire duration.

### Data Analysis

Experimental data (OD_600_ and fluorescence) were processed and the mean and standard deviation were plotted using Origin 2025 software.

## AUTHOR INFORMATION

### Notes

The author declares no financial interest.

## ACKNOWLEDGMENT

We gratefully acknowledge Stanford University and the Department of Mechanical Engineering for their support throughout this work. We also thank the Department of Bioengineering for providing valuable resources and facilities. This research was supported in part by the Burroughs Wellcome Fund. We are thankful to our lab mates in the Mayalu Lab for their insightful discussions, technical assistance, and encouragement during the course of this study.

